# Validation and automation of a high-throughput multi-targeted method for semi-quantification of endogenous metabolites from different biological matrices using tandem mass spectrometry

**DOI:** 10.1101/352468

**Authors:** Jatin Nandania, Gopal Peddinti, Alberto Pessia, Meri Kokkonen, Vidya Velagapudi

## Abstract

The use of metabolomics profiling to understand metabolism under different physiological states has increased in recent years, which created the need for robust analytical platforms. Here, we present a validated method for targeted and semi-quantitative analysis of 102 polar metabolites that covers major metabolic pathways from 24 classes in a single 17.5-min assay. The method has been optimized for a wide range of biological matrices from various organisms, and involves automated sample preparation, and data processing using in-house developed R package. To ensure reliability, the method was validated for accuracy, precision, selectivity, specificity, linearity, recovery, and stability according to European Medicines Agency guidelines. We demonstrated excellent repeatability of the retention times (CV<4%), calibration curves (R^2^≥0.980) in their respective wide dynamic concentration ranges (CV<3%), and concentrations (CV<25%) of quality control samples interspersed within 25 batches analyzed over a period of one-year. The robustness was demonstrated through high correlation between metabolite concentrations measured using our method and NIST reference values (R^2^=0.967), including cross-platform comparability against the BIOCRATES AbsoluteIDQp180 kit (R^2^=0.975) and NMR analyses (R^2^=0.884). We have shown that our method can be successfully applied in many biomedical research fields and clinical trials, including epidemiological studies for biomarker discovery. In summary, a thorough validation demonstrated that our method is reproducible, robust, reliable, and suitable for metabolomics studies.

## 1. Introduction

Metabolomics has a great influence on many disciplines, as metabolites are intermediates or end products of cellular functions. Hence, metabolomics can be used as a powerful tool to generate data for understanding, diagnosing, and managing different pathophysiological conditions. It is therefore essential to be able to identify and measure metabolites from different biological matrices [1]. Although global metabolomics has been widely used in discovery studies for understanding cellular responses to normal and abnormal biological conditions, targeted metabolomics has more advantages for addressing biological questions in a more hypothesis-driven manner than global untargeted metabolomics [2]. Furthermore, targeted metabolomics can quantify metabolites that are low in abundance, which are difficult to assess using an untargeted approach.

An appropriate sample pretreatment is required to obtain reproducible and high quality quantitative data in targeted metabolomics. However, metabolites are present in a wide dynamic range with great diversity in physicochemical properties in the biological matrices [3]. Recent advancements in extraction techniques and automated approaches for sample preparation have partially satisfied the demands of targeted metabolomics. However, there are still many outstanding challenges, such as the matrix effect and laboratory-to-laboratory variation associated with sample preparation. Hence, standardization of sample preparation is a fundamental requirement in metabolomics studies [4].

Secondly, robust analytical methodology is required for accurate quantification of metabolites with good reproducibility over an extended period of time [5]. Molecular diversity is a major problem that hinders the separation of all pre-selected metabolites in a single chromatographic run and the detection of all separated metabolites with minimum technical variation [6]. Tandem mass spectrometry (MS) is a technique used predominantly due to its high sensitivity and high throughput for the detection of metabolites. A combination of MS and separation techniques is used to increase the sensitivity and reliability of analytical methods for the analysis of metabolites from complex biological matrices [7]. In addition, latest developments in triple quadrupole instrumentation strengthened the possibilities to develop multi-analyte methods in a single injection that yield reliable and quantitative data [8].

Even though liquid chromatography-mass spectrometry (LC-MS) is a method of choice in targeted metabolomics, obtaining accurate quantification and long-term data reproducibility remains an analytical challenge. This is due to limitations such as matrix effect, MS performance drift, and LC column contamination and aging [9]. Finally, although instrument vendor software for data processing provides some crucial functions such as peak integration (rendering the data from high-throughput metabolomics experiments into numerical values that represent metabolite concentrations), the lack of automation in downstream data processing and quality control remains a major bottleneck in high-throughput analyses. Thus, an efficient pipeline is necessary to enable rapid, accurate, and standardized processing of these data. Such a pipeline should facilitate automated analyses, while at the same time allowing the user to fine-tune the parameters for accurate data processing. Taken together, there is a need for development and validation of standardized, robust, and quantitative methods for large-scale targeted metabolomics studies in a high-throughput manner to minimize bias associated with sample preparation and the analytical technique used.

We have previously developed a robust, reproducible, and high-throughput targeted method for measuring 102 polar metabolites from various biological classes semi-quantitatively in a single injection [10,11]. The metabolites were selected according to the following criteria: have important roles in many biological processes, are known biomarkers in several diseases and technical feasibility for developing an analytical method that covers all the metabolites in a single assay. The selected metabolites come from 24 different classes (Table 1), covering a wide range of metabolic pathways. We created an in-house metabolite database by manually curating all the available information (i.e., names, HMDB, PUBCHEM, KEGG Ids, chemical properties, reported normal and abnormal concentration ranges, links to their structures) from the Human Metabolome Database (HMDB).

The selected metabolites were separated by using Hydrophilic Interaction Liquid Chromatography (HILIC) and measured using triple quadrupole mass spectrometry (MS). Briefly,after addition of an internal standards working solution, biofluid samples were extracted with 1% formic acid in 99% acetonitrile. Tissue samples were extracted using homogenization in two steps. Extraction was first performed with 1% formic acid in 99% acetonitrile; the second extraction used 80/20% ACN/H_2_O+1% formic acid by protein precipitation followed by filtration through Ostro plates. After this step, 5 μL of filtered sample extract was injected into an Acquity UPLC system coupled to a Xevo^®^ TQ-S triple quadrupole mass spectrometer (Waters Corporation, Milford, MA, USA), which was operated in both positive and negative polarities with a polarity switching time of 20 msec for metabolite separation and quantification. Multiple Reaction Monitoring (MRM) acquisition mode was selected for metabolite quantification. A detailed method description and instrument parameters are provided elsewhere [10,11]. Our method involves automated sample preparation for biofluid samples and minimal manual steps for non-biofluid samples. This reduces the analysis time and inter-batch variation and minimizes human error. Semi-quantification of the metabolites was performed using external calibration curves (R^2^≥0.980) in their respective wide dynamic concentration ranges and 12 labeled internal standards were used for most of the metabolites to minimize matrix effects.

Several analytical methods have also been developed for semi-quantitative measurement of large number of metabolites in a single run [12–16]. Li et al [12] measured 610 metabolites from 60 biochemical pathways. However, their method requires approximately 40 minutes to separate all the metabolites (with poor resolution for some metabolites). Furthermore, their method was not applicable to all biological matrices. Similarly, Wei et al [15] detected approximately 200 metabolites in 10 minutes from plasma. However, their method lacks information on validation and for other biological matrices. Yuan et al [16] also provided a protocol for detection of approximately 250 metabolites in a single method. However, they did not use an automated platform and a drying step in their method increases the analysis time of the method overall. Our method has clear advantages compared to other published methods, including shorter analysis time, high throughput, automation, and semi-quantitation using individual external 11-point calibration curves along with 12 labeled internal standards. Furthermore, the method has been used in many biomedical and clinical studies for biomarker discovery [10,11,17–37].

The primary objective of this work is to show the robustness of our method through a thorough validation of our previously developed analytical method according to European Medicines Agency (EMA) guidelines. We also demonstrate automation of tedious and manual data processing tasks in high-throughput metabolomics analyses using an in-house developed R package. The R package automates various corrections and normalization steps to convert the raw peak area data to molecular concentrations for each compound in each sample and also provides quality evaluation of the data and reduces the manual work load significantly.

## 2. Materials and Methods

### 2.1 Chemicals and reagents

All metabolite standards were purchased from Sigma-Aldrich (St. Louis, MO, USA). Internal standards were ordered from Cambridge Isotope Laboratory. Inc. (Tewksbury, MA, USA). LC-MS-grade solvents, 2-proponol, acetonitrile, and methanol (HiPerSolv) were obtained from VWR International (Helsinki, Finland). Analytical-grade chemicals (formic acid, ammonium formate, and ammonium hydroxide) were obtained from Sigma-Aldrich. Deionized water (18 MΩ.cm at 25°C) used for solution preparation was made using a Milli-Q water purification system (Bamstead EASYpure RoDi ultrapure water purification system, Thermo scientific, Ohio, USA). Mouse tissues, including heart, liver, brain, spleen, and muscles were obtained from Innovative Research Laboratory (Novi, MI, USA). Whole blood was obtained from the Finnish Red Cross blood service (Helsinki, Finland), from which serum was prepared during method optimization and validation. Cell samples were provided by our research collaborators. NIST Standard reference material (SRM) 1950 plasma was purchased from Sigma-Aldrich (Gillingham, UK).

### 2.2 Metabolite extraction protocol and instrumentation

All metabolites were extracted, separated with HILIC chromatography (Acquity BEH amide, 2.1 X 100 mm, 1.7μ), and analyzed with a Waters Xevo TQ-S mass spectrometer using our previously published protocol [11]. The protocol for tissues and adherent cells was optimized for better recovery and chromatography and to cover a wide range of tissue and cell types with a single protocol. For tissue sample extractions, 90/10% ACN/H2O + 1% formic acid was used instead of 80/20% ACN/H2O+ 1% formic acid during the second step of extraction. Additionally, during cell pellet sample extraction, 80/20% ACN/H2O+1% formic acid was replaced with 90/10% ACN/H2O + 1% formic acid. After optimization, we used the tissues protocol for analysis of various biological matrices, such as heart, liver, placenta, brain, muscles, spleen, *C. elegans, Drosophila* larvae, dental carries, dried blood spots, and fecal samples. The cell pellet protocol was used for all types of adherent cells and *E. coli* and *S. cerevisiae* samples. The biofluid protocol was used for all types of biofluids, such as blood, plasma, serum, cell culture supernatant, CSF, and urine.

### 2.3 Method Validation

Validation of the method was performed to verify various parameters and the reliability of the developed method for analysis of a large number of samples. The method was validated according to EMA guidelines for bioanalytical method validation in terms of selectivity, specificity, linearity, accuracy, precision, extraction recovery, matrix effect, and stability [38]. In addition, we used pooled healthy human serum samples as internal quality control (QC) samples in all studies to correct signal drift during sample runs and to improve confidence in the statistical data. QC samples at high, medium and low (for serum) or high and low concentration levels (for tissues) were prepared by spiking a mixed standard solution in their respective homogenized biological matrices to perform all the method validation experiments. We performed validation for commonly used biological samples in metabolomics analyses, such as biofluid (serum), tissue (liver, brain and spleen), and cell samples. An aqueous calibration curve was used to calculate the concentration values during the method validation. The instrument performance for response reproducibility and sensitivity was always verified by six consecutive injections of medium concentration solution at the start of any experiment.

#### 2.3.1 Selectivity and specificity

The selectivity and specificity for each metabolite were investigated using serum-spiked samples (N=6) with known amount of standard. Chromatographic interferences from other endogenous compounds of the biological matrix at the retention time of the target analyte for a particular metabolite were verified. The chromatographic peaks from spiked samples were compared with the standards by the retention times and if required from their respective MRM spectra.

#### 2.3.2 Linearity, accuracy, and precision

To assess the linearity, accuracy, and precision, six replicates of spiked QC samples at high, medium, and low concentrations along with calibration curve were injected on three separate days. Calibration curve standards of 11 points were prepared via serial dilution. The curve was plotted by using the peak area response ratios (standard/labeled standard) versus the concentrations of the individual metabolites. The calibration curve was constructed using the regression equations (linear and quadratic) by applying appropriate weighing factor and by transforming the axis (both instrument response and theoretical concentration) into logarithmic or square root function. The accuracy was calculated as measured value divided by the nominal value at each concentration level of the calibration curve standards in all three batches. Inter-and intra-batch variability was calculated by measuring coefficient of variation (%CV) at each QC concentration level.

#### 2.3.3 Recovery and matrix effect

The recovery efficiencies for each analyzed metabolite was determined by comparing analytical results from QC samples spiked with standards mixture before and after extraction using different concentrations. The spike concentrations covered the calibration range. The matrix effect (percentage of ion suppression or enhancement of the MS signal) was determined by comparing the analytical response of QC samples that were spiked after extraction with the analytical response of aqueous spiked samples (diluent spiked with respective concentrations of QCs). Since there were endogenous metabolites, we subtracted the endogenous concentrations from the samples that were spiked. This experiment was performed using six QC replicates.

#### 2.3.4 Stability of the metabolites

Wet extract, freeze-thaw, and stock solution stability for all metabolites were determined to check the integrity of the analytes in solvents and in QC samples at different conditions. To determine the wet extract stability, six replicates of extracted QC samples were kept in the auto-sampler at 5°C. The same samples in the same sequence were reinjected with freshly extracted QC samples and the results were compared.

Freeze-thaw stability was evaluated up to three cycles by freezing and thawing the spiked QC samples stored at −80°C and comparing the concentrations against the freshly thawed and spiked QC samples.

Long-term stock solution stability for metabolite stock solutions and intermediate solutions were checked by comparing the mean peak area of freshly prepared solutions with stored solutions at 4°C. All stability experiments were performed with six replicates of QCs.

#### 2.3.5 Sample carry-over

Sample carry-over was evaluated by injecting the highest standard concentration (ULOQ) of the metabolites in the calibration curve followed by a series of blank injections and lowest standard concentration (LLOQ). The blank samples were evaluated for any signal at the retention time of particular metabolites and signal intensities of the blank samples were compared with the LLOQ samples. The acceptance criteria for carry-over was set at 20% of the peak area corresponding to the LLOQ level as per the EMA guidelines for bioanalysis.

#### 2.3.6 Quality control samples

Internal QC samples were prepared after separating serum from pooled healthy human blood samples. A volume of 350 μL of serum was aliquoted and stored at −80°C after providing a lot number and QC number. The concentration of QC samples that were incorporated in batches during the metabolomics studies were calculated for all the metabolites along with the experimental samples. Average concentrations (μmol/L) and %CV of the QC samples were calculated for each metabolite. The data were saved along with QC lot numbers, batch name, and run date. The QC data were collected from six different lots for a period of 5.5 years (N=539 replicates). An internal QC database has been maintained and used for quality checks.

#### 2.3.7 Comparison with reference material

To evaluate the performance of our semi-quantitative method, commercially available standard reference plasma (NIST SRM 1950) [39] was analyzed using our method (N=8 replicates). The concentration values from the matched 17 metabolites were compared with the given standard reference values.

#### 2.3.8 Cross-platform comparison

To further evaluate the robustness and performance of our method, we performed a cross-platform comparison using two completely different analytical platforms: (1) the commercially available AbsoluteIDQ p180 targeted metabolomics assay kit using LC-MS/MS and (2) a nuclear magnetic resonance (NMR) platform. We sent our internal QC samples to the BIOCRATES Life Sciences AG (Innsbruck, Austria) (N=3 replicates) and to the NMR Metabolomics Laboratory, School of Pharmacy, University of Eastern Finland (Kuopio, Finland) (N=3 replicates). Our QC samples were extracted and analyzed as described previously for the AbsoluteIDQ p180 kit [40] and for the NMR analysis of small molecules [41]. We compared these results with the results obtained from our method (N=4-5 replicates).

### 2.4 Statistical analyses

To estimate the median concentration values (μmol/L) of the metabolites from QC samples (N=539), we fitted a linear mixed model with the MCMCglmm R package [42] using an expanded parameter formulation and default settings. Observed data was assumed to be log-normally distributed and corrected for the six different QC lots. Credibility intervals (95%) of median concentration values were computed from 20 000 samples of the posterior distribution. Error bars shown in the scatter plots are 95% confidence intervals, while the coefficient of determination R^2^ refers to the simple linear regression between the two plotted variables. Coefficient of variation (CV) percentages were calculated as a measure of variability. To automate the downstream processing of the data produced by the instrument vendor software (TargetLynx), we built a data processing package called “Unlynx” in R statistical programming language.

### 2.5 Automated data processing

The “Unlynx” package parses the output of TargetLynx software (i.e., raw data containing the concentration values in PPB units) and produces a processed dataset in an Excel spreadsheet after performing a series of preprocessing operations.

The preprocessing steps included the following:

i. Molecular weight normalization, in which the ppb values are normalized by the molecular weight of each compound, thereby converting the data from ppb units to μmoles.
ii. Process efficiency correction for the semi-quantification of metabolites without internal standards.
iii. Normalization using dilution factor for specific sample type if dilution was needed.
iv. Cell number normalization (for cell samples) to convert the concentration values per million cells.
v. Calculation of mean, standard deviation, and relative standard deviation (RSD) of molecular concentrations (resulting from the previous steps) for each phenotypic group.
vi. Outlier detection in each phenotypic group; if the concentration value of a compound in a sample is more than one or two standard deviation (SD) away from the mean of the phenotypic group, then it is marked as an outlier in the Excel data set in two different colors.
vii. QC check by comparing the RSD of QC samples in the current dataset against the internal database of QC sample RSDs (based on inter-day RSDs recorded over one year).

## 3. Results and Discussions

### 3.1 Extraction method optimization

The primary objective of this work was to optimize and validate our previously published protocol for different types of biological matrices. For tissue samples (placenta, liver, heart, brain, spleen, and muscles), the sample volumes of the tissues and extraction solvent volumes were optimized to fit the concentrations of most of the metabolites within the linearity of calibration curve for reliable results. We observed that most of the metabolites can be semi-quantified within the calibration curve range with 20±5mg of sample weight.

Furthermore, we optimized the protocol with extraction solvent for tissues and adherent cells. Some of the metabolites (in particular inositol, GABA, asymmetric dimethylarginine, symmetric dimethylarginine, spermidine, ribose-5-phosphate, and orotic acid) had poor separation and irreproducible chromatography. Interference of isobaric compounds with other metabolites was also observed due to poor separation. Thus, different compositions of the extraction solvent were assessed to achieve the acceptable chromatography. We observed that modification of acetonitrile content from 80% to 90% and applying longer equilibration time for the HILIC column yielded acceptable chromatography and also good separation for most of the metabolites.

We also optimized the extraction protocol for different sample types from various organisms, such as tissue types (adipose, endometrium, testicles, lung), fecal samples, dried blood samples, dental carries, biofilm, extracellular vesicles, mitochondrial isolates, drosophila, *C. elegans, E. coli*, and *S. cerevesiae* samples (Figure 1).

**Figure 1.**
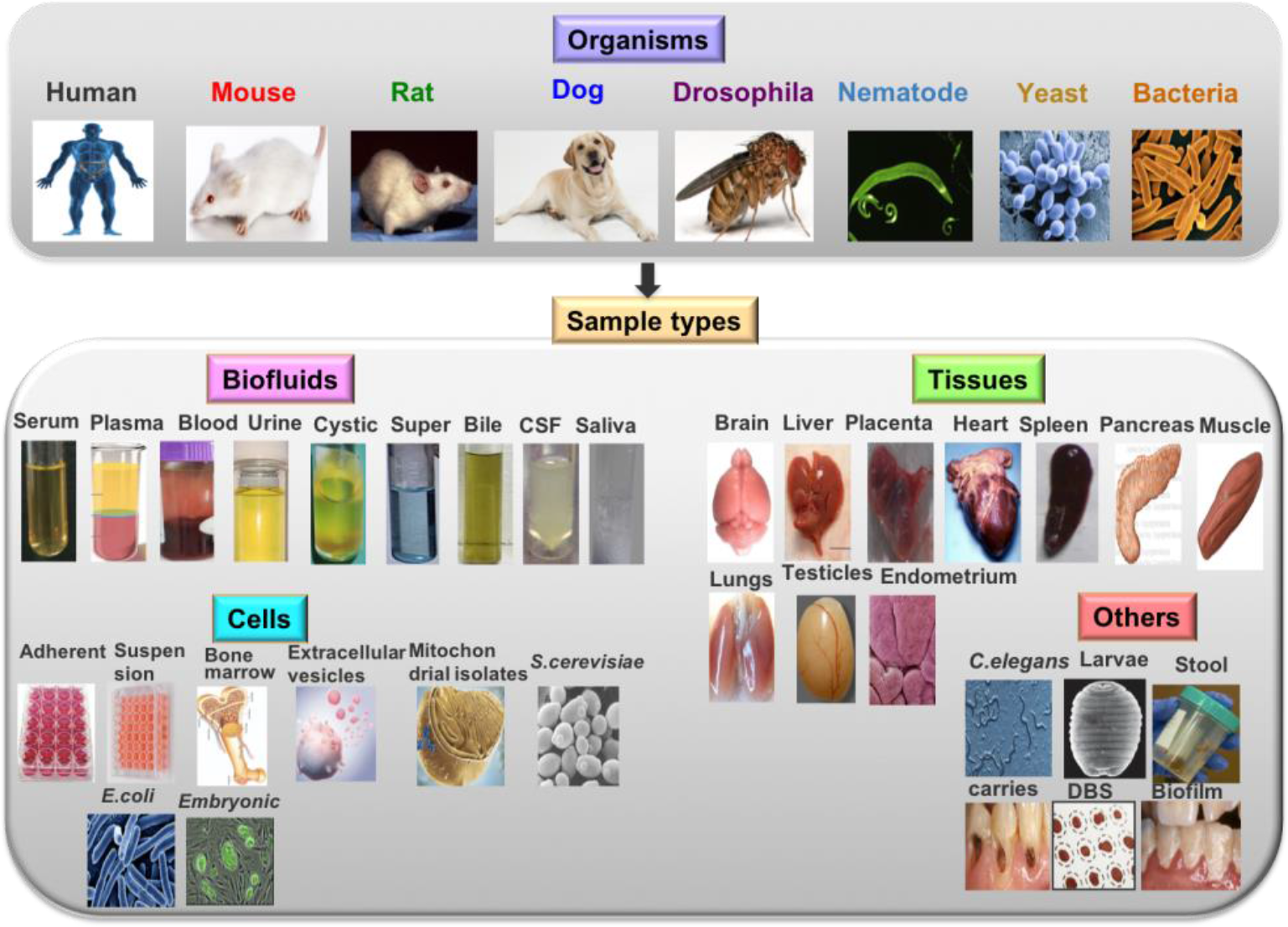
Illustration of different sample types from various organisms that have been used for extraction method optimization.

### 3.2 Method Validation

#### 3.2.1 Selectivity and specificity

There were no significant interference peaks from the matrix components in their respective retention time windows, indicating the selectivity of the metabolites in our method. We repeated the injections for five times from all different serum samples and confirmed that the peak eluted was only from the target analyte, indicating that they are specific to their corresponding MRM transitions (Figure 2).

**Figure 2.**
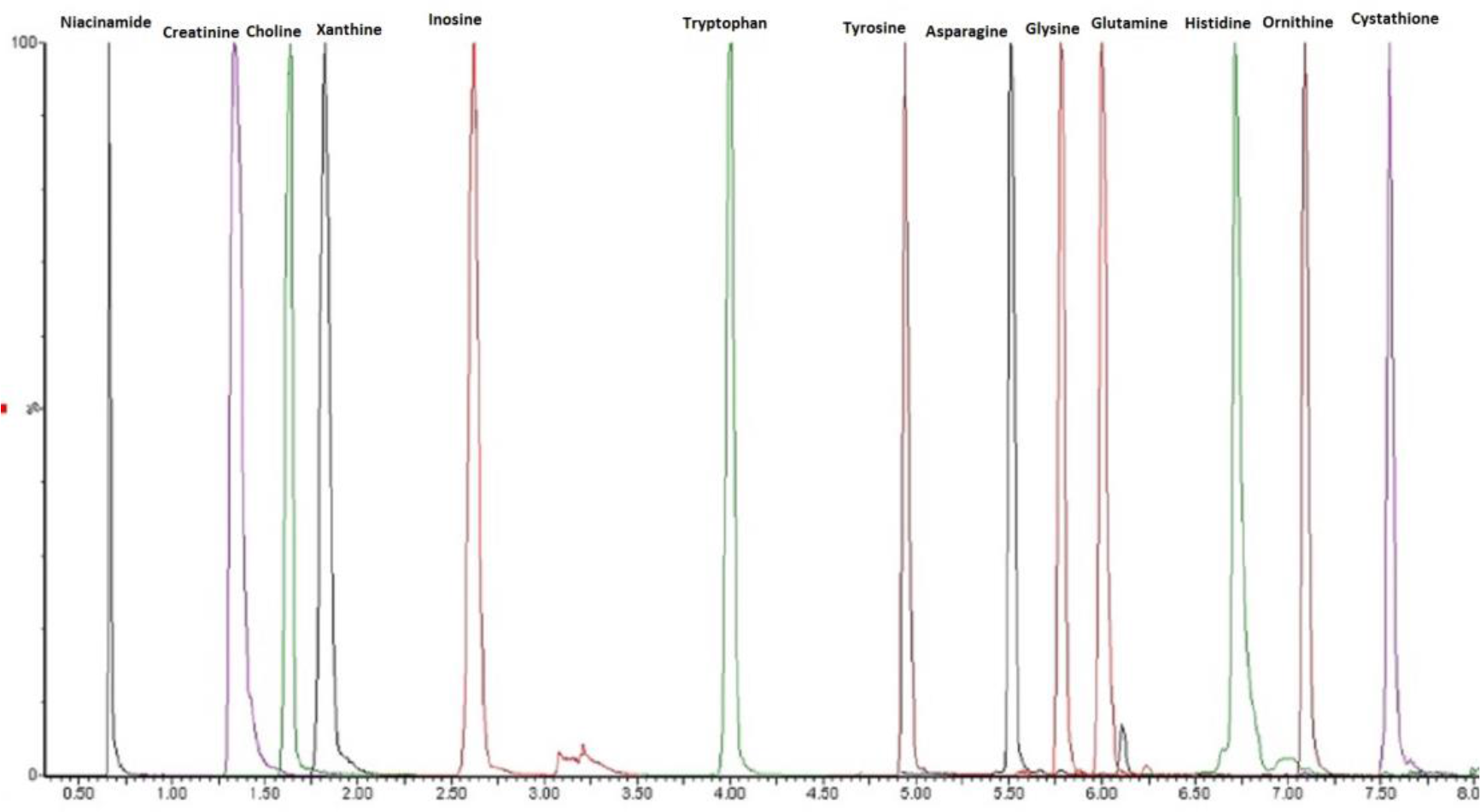
Representative chromatographic peaks of selected metabolites at their respective retention times in the QC serum sample.

#### 3.2.2 Linearity, accuracy, and precision

To cover a broad concentration range, the linear or quadratic models were used and the variables from X and Y axes were logarithm or squareroot transformed to fit the calibration data [43]. The coefficient of determination (R^2^) value for each metabolite was greater than 0.980 at their respective concentration range, except for some metabolites such as aspartate, uracil, 2-deoxyuridine sucrose, and chenodeoxycholic acid (likely due to their broad peak shapes and poor recovery at lower concentration, Table S1).

Concentration precision for QC samples was calculated by measuring %CV at high, medium, and low concentration level of QCs (N=6 replicates). In general, intra-and inter-day precision (CV) values were within 15% for most of the metabolites except acetoacetic acid, folic acid, sucrose, homoserine, 2-deoxyuridine, and cholic acid (Figure 3). However, at low concentrations, more than 20% CV were observed for NAD and myo-inositol. This might be due to the low recoveries of these compounds.

**Figure 3.**
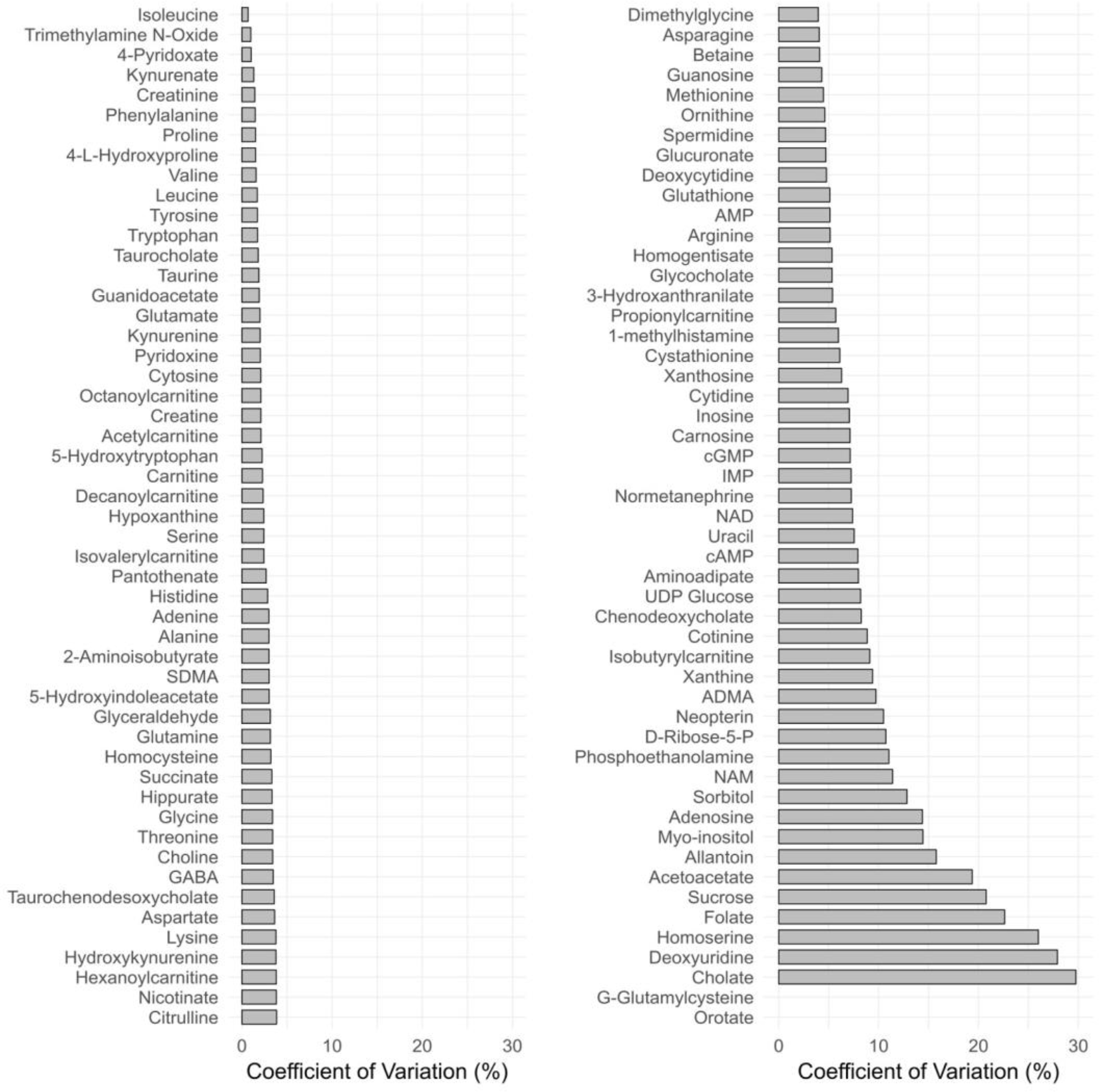
Inter-batch variation of high concentrations of spiked QC serum samples (N=6) analyzed on 3 different days. Data not shown for g-glutamylcysteine and orotic acid (CV>30%).

#### 3.2.3 Recovery and matrix effect

For most of the metabolites, recoveries were found to be between 50% to 120% with good repeatability at all three concentrations levels (low, medium, and high) in both biofluid (serum) and tissues (brain, liver, and spleen). However, compounds such as UDP-glucose, IMP, cGMP, D-ribose 5-phosphate, NAD, AMP, homocysteine, carnosine, and glutathione had recoveries less than 30% in serum. However, CV of recoveries at low, medium, and high concentration levels were within 25% except for cGMP in serum. Metabolites such as 1-methylhistamine, aspartate, glutamine, adenosine, and glutathione had recoveries over 120%. Histidine, ornithine, cystathionine, 3-OH-DL-kynurenine, carnosine, AMP, NAD, cGMP, IMP, and UDP-glucose had less than 30% recovery in some tissues types, indicating matrix effect or degradation (Figure 4). However, CV of repeatability at all concentration levels were within 15% except for 1-methylhistamine, aspartate, glutamine, glutathione, NAD, and UDP-glucose in some tissues.

**Figure 4.**
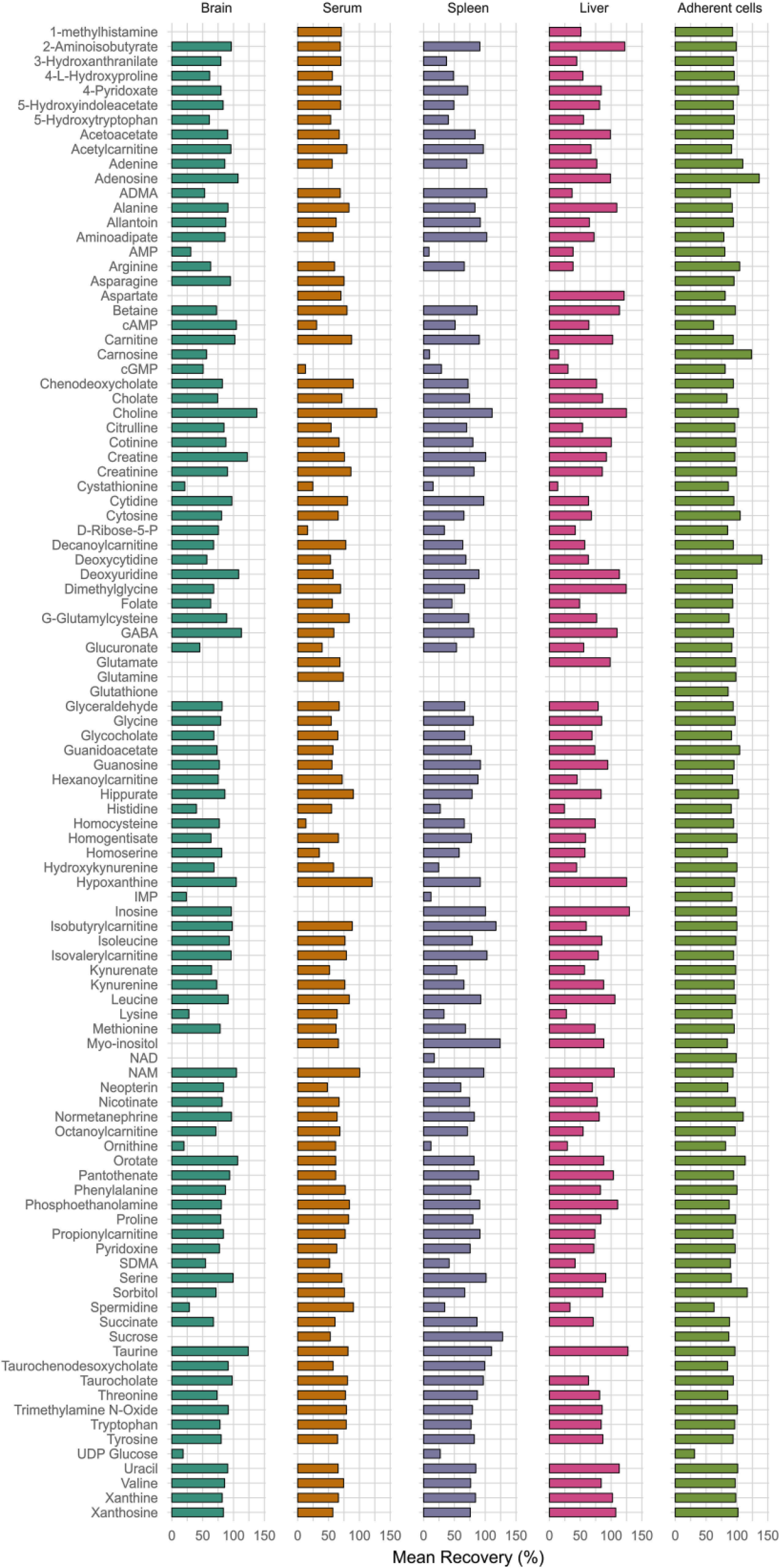
Percentage mean recoveries for all metabolites at low, medium, and high concentration levels of QCs (N=6 at each level) spiked in serum, brain, spleen, liver, and adherent cells. Data not shown for metabolites with poor chromatography and irreproducible results at different concentration levels.

The matrix effect values were observed to be within the range of 0.6 to 1.8 (below 1 indicates ion suppression and above 1 indicates ion enhancement) for the metabolites in serum and tissues. The challenge of the matrix effect can be overcome by having individual isotope-labeled internal standards for each individual compound for true quantification. However, this is not practically possible for high-throughput metabolomics analyses, where usually the aim is relative comparison of cases *vs* controls. This is due to high costs and also because not all internal standards are commercially available. In our method, we selected 12 labeled internal standards (Table S1), which represent chemically similar classes for optimal correction. This is because the matrix effect was expected to be the same for an analyte and its labeled isotope analogue. The process efficiency percentages were calculated for the metabolites without internal standards. The analyte concentrations determined through the external calibration were divided with the total process efficiency values to correct the concentration values of the analyte in the given biological sample. Also, the repeatability of matrix effect in terms of CV was less than 25% for most of the compounds. Reliable measurements are accordingly possible.

#### 3.2.4 Stability

Endogenous metabolites are not stable due to degradation or conversion reactions. Hence, the stabilities of all metabolites were assessed under different conditions. For wet extract stability, approximately 90% of the metabolites were stable (stabilities range between 85% and 115%) for 35 hours at 5°C in the auto-sampler (Figure 5A).

**Figure 5.**
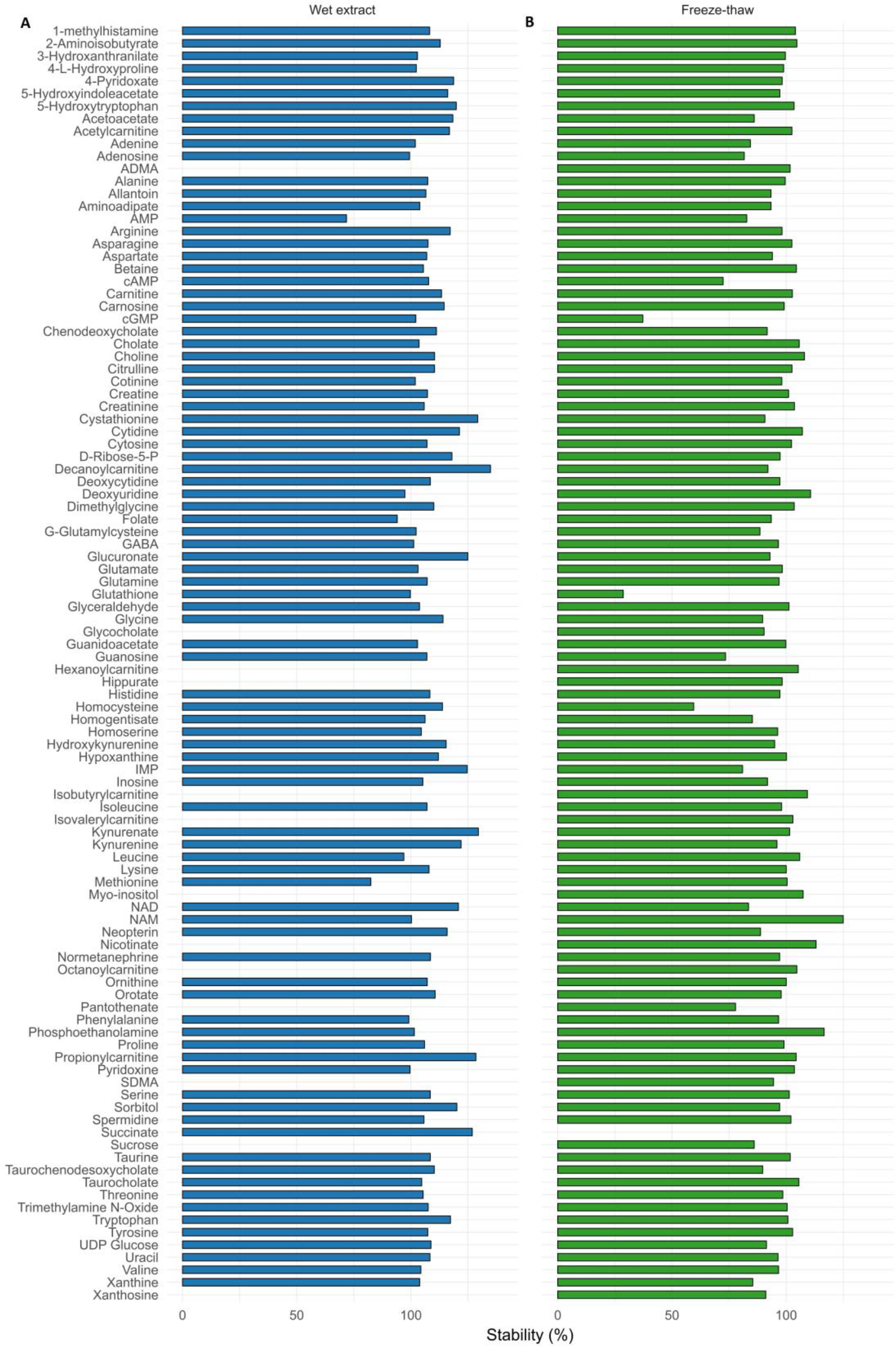
Percentage mean stability was calculated for three different concentrations (low, medium, and high) spiked in QC serum (N=6 at each level) during (A) wet extract and (B) freeze-thaw stability for all the metabolites.

For freeze and thaw cycle stability, most of the metabolites were stable even after three freeze and thaw cycles, with the exception of cGMP, succinate, glutathione, and homocysteine (stability below 30%, Figure 5B). This information is particularly important for clinical studies, where samples are often thawed once or twice.

To determine the stability of working solutions, we started evaluating the stability from intermediate solutions for all the metabolites. Most of the metabolites were stable for 245 days at intermediate concentration when stored at 4°C. However, 16 metabolites had low stability at intermediate concentration; stabilities were thus determined at stock-level concentration for these compounds. We observed that the stock solutions were stable for 56 days except taurocholic acid, sucrose, UDP-glucose, and glutamine (Figure 6). Hence, these stock solutions were freshly prepared during the analysis. The stability of internal standards solutions was also assessed and they were stable for one year.

**Figure 6.**
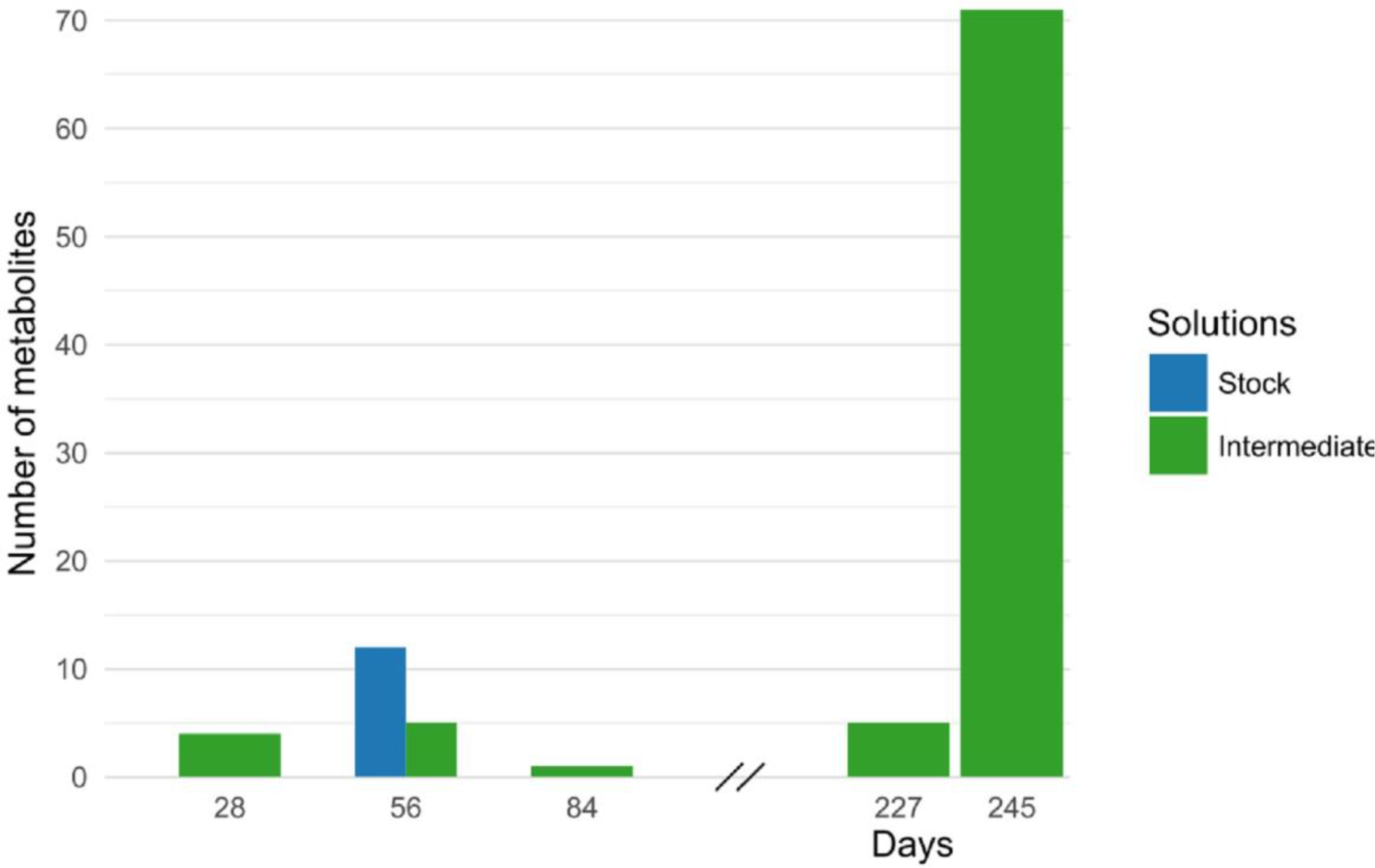
Bar graph representing stability of stock solutions and intermediate solutions of metabolites that were used to freshly prepare calibration curve standards during the analysis

#### 3.2.5 Sample carryover

In general, for majority of the analyte MRM channels neither peak nor any interference in the blank samples was detected after injection of metabolite standard with high concentration. For compounds such as spermidine, succinate, AMP, and IMP, carryover eluted constantly even after washing but was not significantly high. Other than these compounds, we can conclude that the column, needle, syringe, and seal washes were sufficient to avoid any inter-sample carryover.

#### 3.2.6 Reproducibility

To ensure good quality of the data, internal QC samples were incorporated to a batch of samples and run after every tenth experimental sample. QC data were collected from 25 different batches that were performed during various metabolomics studies over a period of 1 year. Mean concentrations and %CV values of QC replicates within each batch were calculated for all the 25 batches. We observed that approximately 80% to 85% of the metabolites were always present within 25% of CV values (Figure 7). The higher %CV values for the remaining metabolites could be partially explained by low abundance in human serum, low recovery, or poor chromatography; these were consistently found to be below LLOQ within the 25 batches.

In addition, %CV values for retention times and R^2^ values of calibration curves for each metabolite in all the 25 batches were calculated to verify the reproducibility. Based on these results, the repeatability was excellent except for a few compounds over a period of 1 year. No drifting effect for the retention times (CV<4%) was observed, and excellent reproducibility was observed for R^2^ values of calibration curves (CV<3%) (Figure 8). On the basis of these results, our method can be considered accurate, reliable, and reproducible.

**Figure 7.**
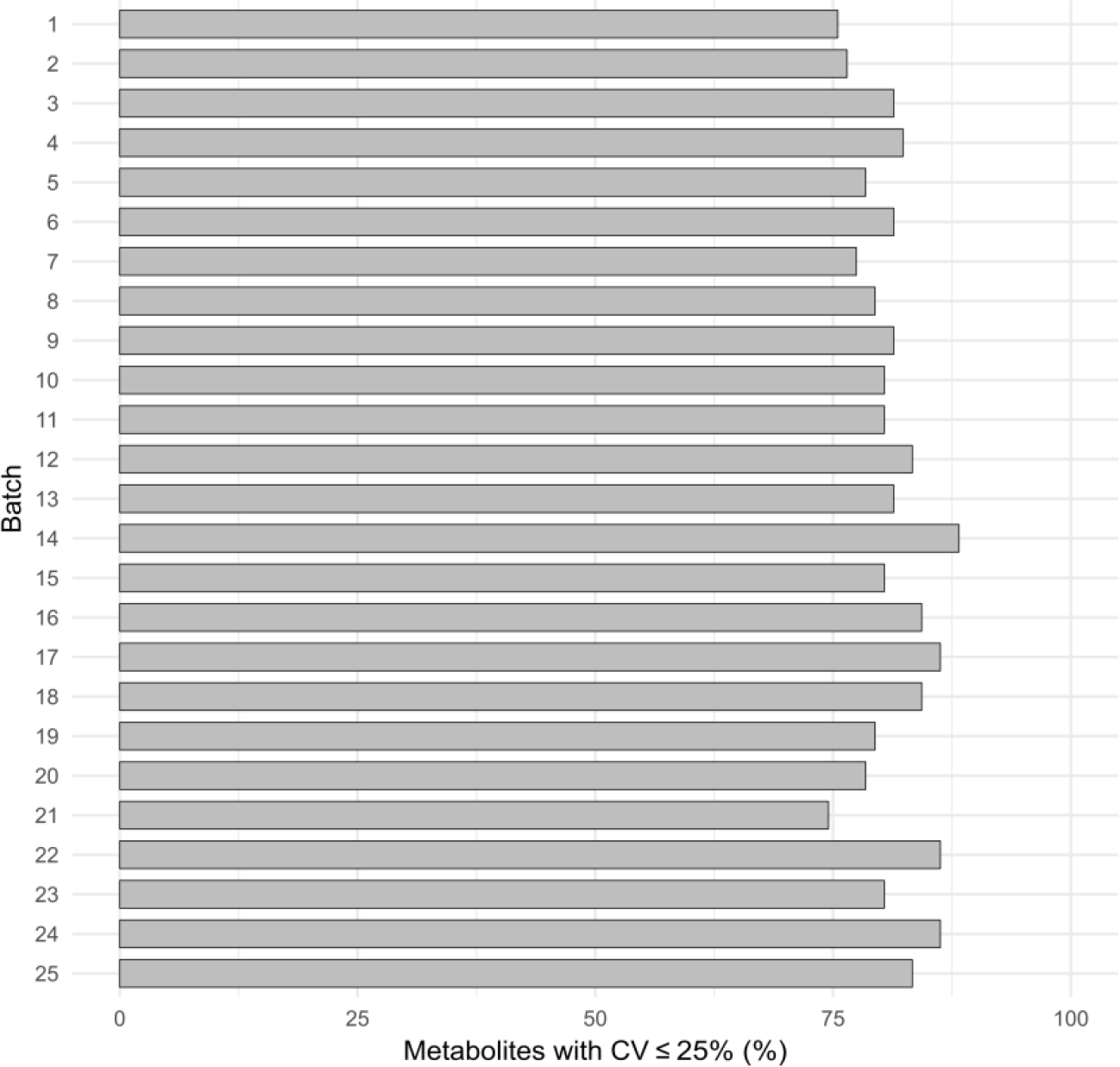
Percentage of metabolites with less than 25% CV values of QC concentrations in 25 different batches analyzed over a period of 1 year.

**Figure 8.**
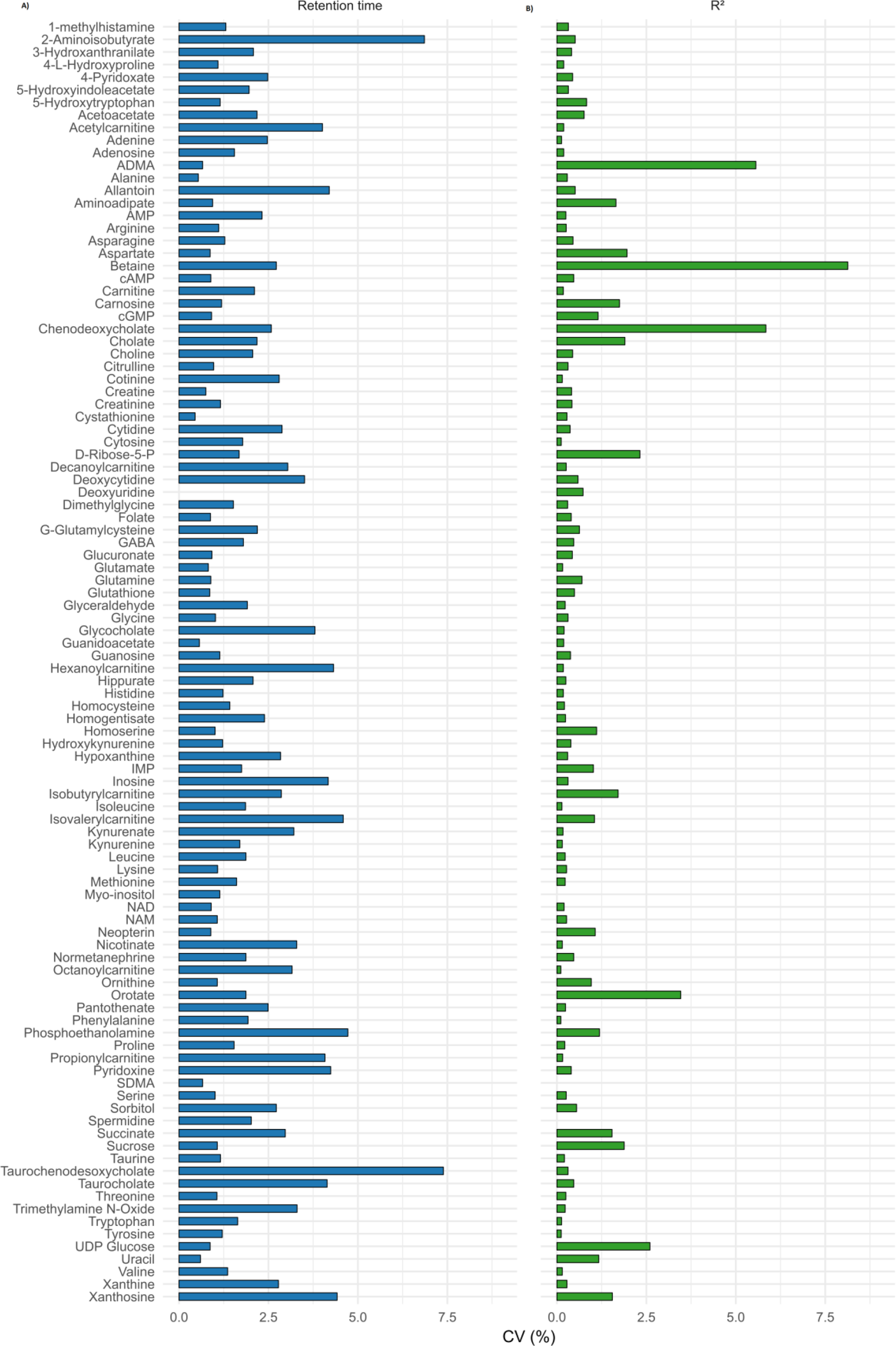
Reproducibility of (A) retention time of respective chromatographic peaks and (B) regression coefficient values (R^2^) from external calibration curve standards for each metabolite analyzed in 25 different batches over a period of 1 year.

#### 3.2.7 Quality management

To obtain reproducible and accurate data, we set up a strict quality management and electronic lab notebook system. To reduce the bias from sample analysis, we always double-randomized the samples (i.e., one before the sample extraction step and one before injecting into the LCMS system across different phenotypes of the samples). For stabilization of response and retention time, we always verified a few runs of highest calibration level 11 before injecting the experimental samples. During the stabilization process, we also verified the chromatography including peak shape, retention time, and response of all the metabolites. Any significant changes in the intensity, peak shape, retention time, and system pressure were thoroughly investigated and corrected by resolving the problems before injection of experimental samples.

To ensure the integrity of LCMS runs, QC samples were run at every tenth experimental sample and a blank sample at every fifth run during all the metabolomics studies within a batch. Furthermore, chromatography and response of QC samples (including chromatography of some metabolites and IS response variation) and blank runs were always verified after completion of the runs and before starting data processing. In case of any abnormality observed for particular samples, those samples were reinjected or reanalyzed. Only after passing these quality checks we proceed further to process the data. This included verifying the accuracy of calibration curve standards, chromatography peak integrations, IS response variation, and verifying LLOQ and ULOQ for each sample for all metabolites within a batch. The high-throughput targeted metabolomics workflow is shown in Figure 9.

**Figure 9.**
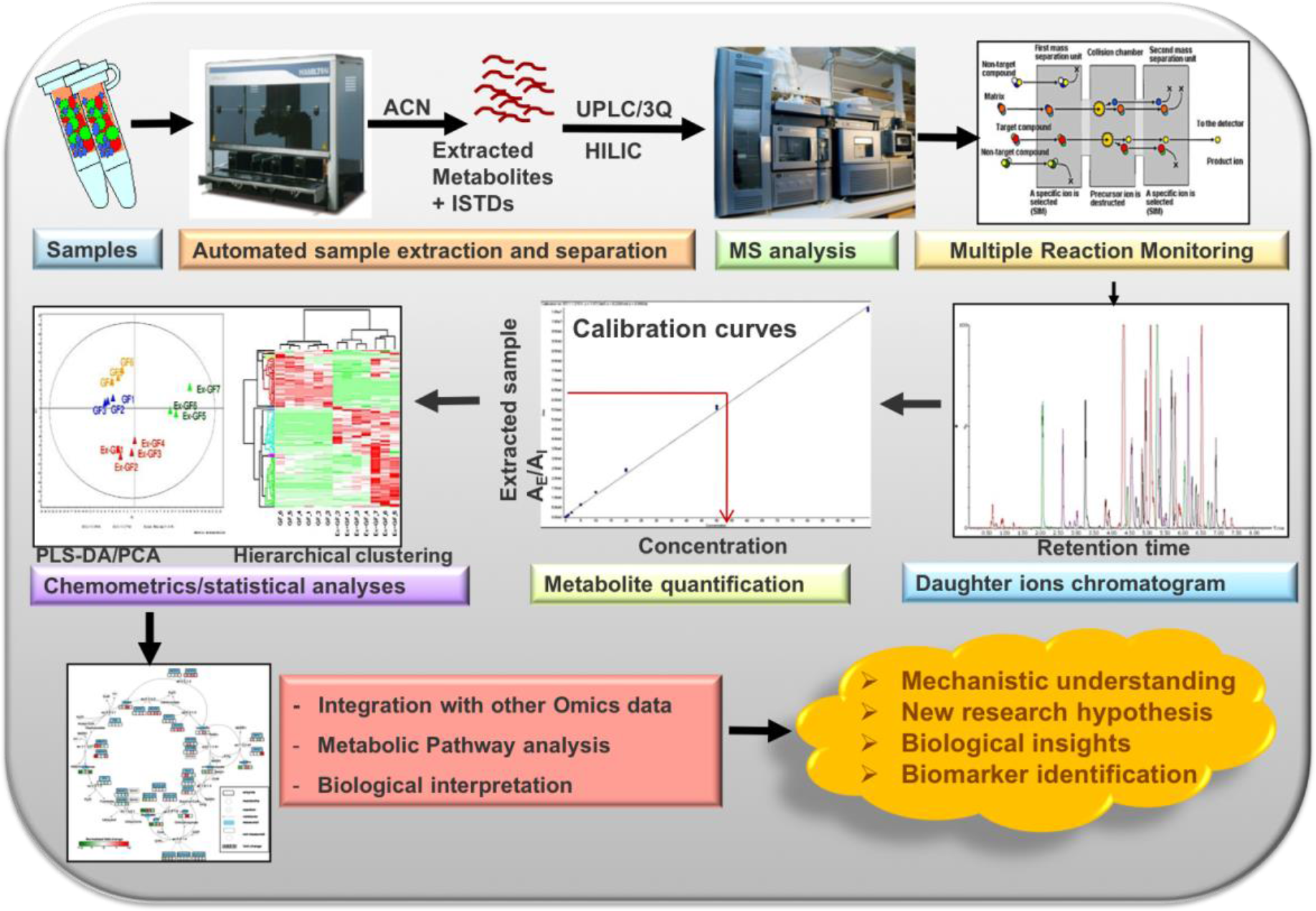
A typical workflow for high-throughput targeted metabolomics analysis.

We collected concentration values (μmol/L) for our QC samples within metabolomics studies conducted over a period of 5.5 years from six different lots (N=539 replicates). The median values of each metabolite together with a 95% credibility interval are presented in Table 1. These represent a reference level for a population of healthy adult individuals.

**Table 1.**
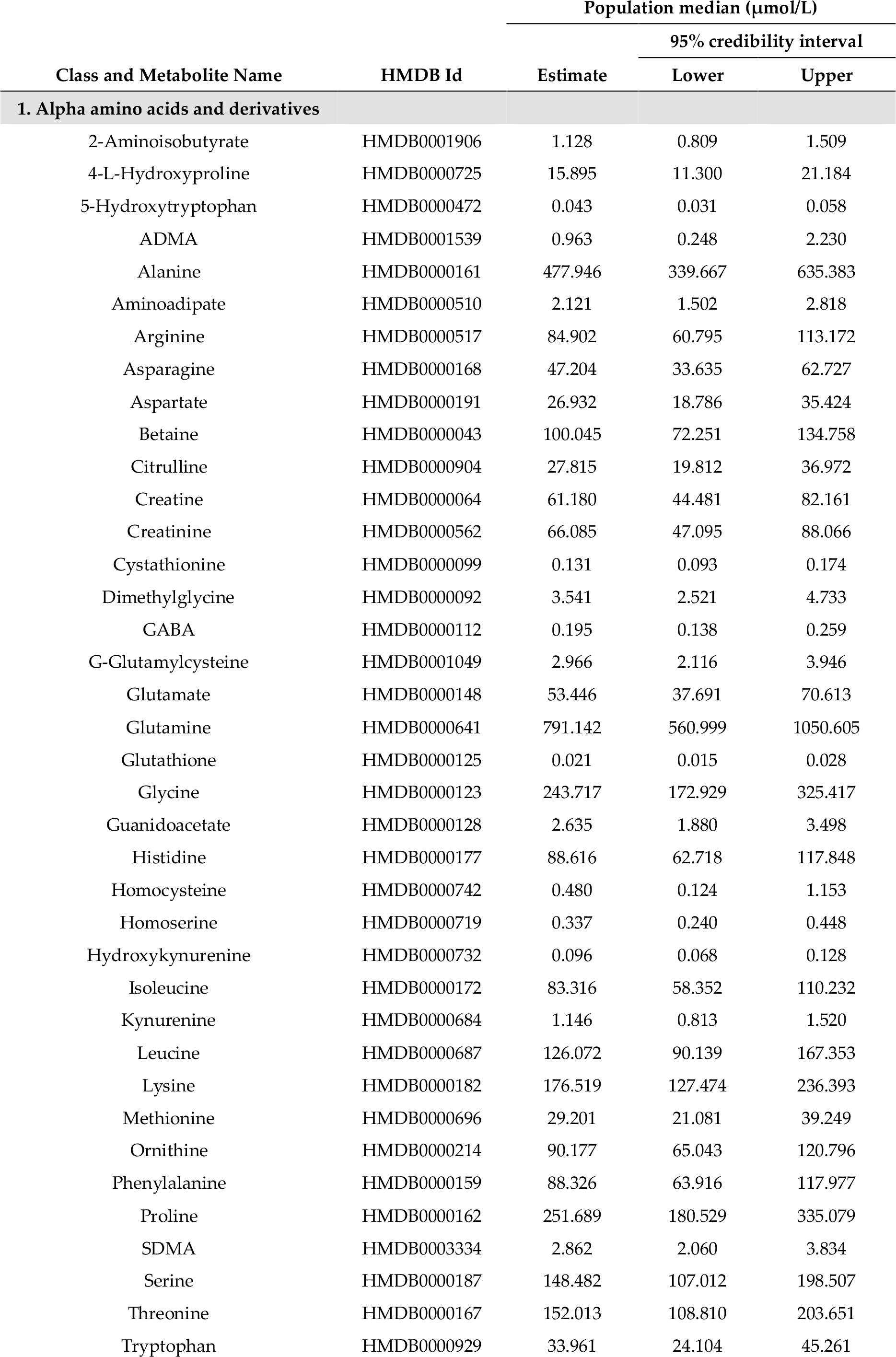
Median concentration levels (μmol/L) measured in pooled healthy adult serum samples.

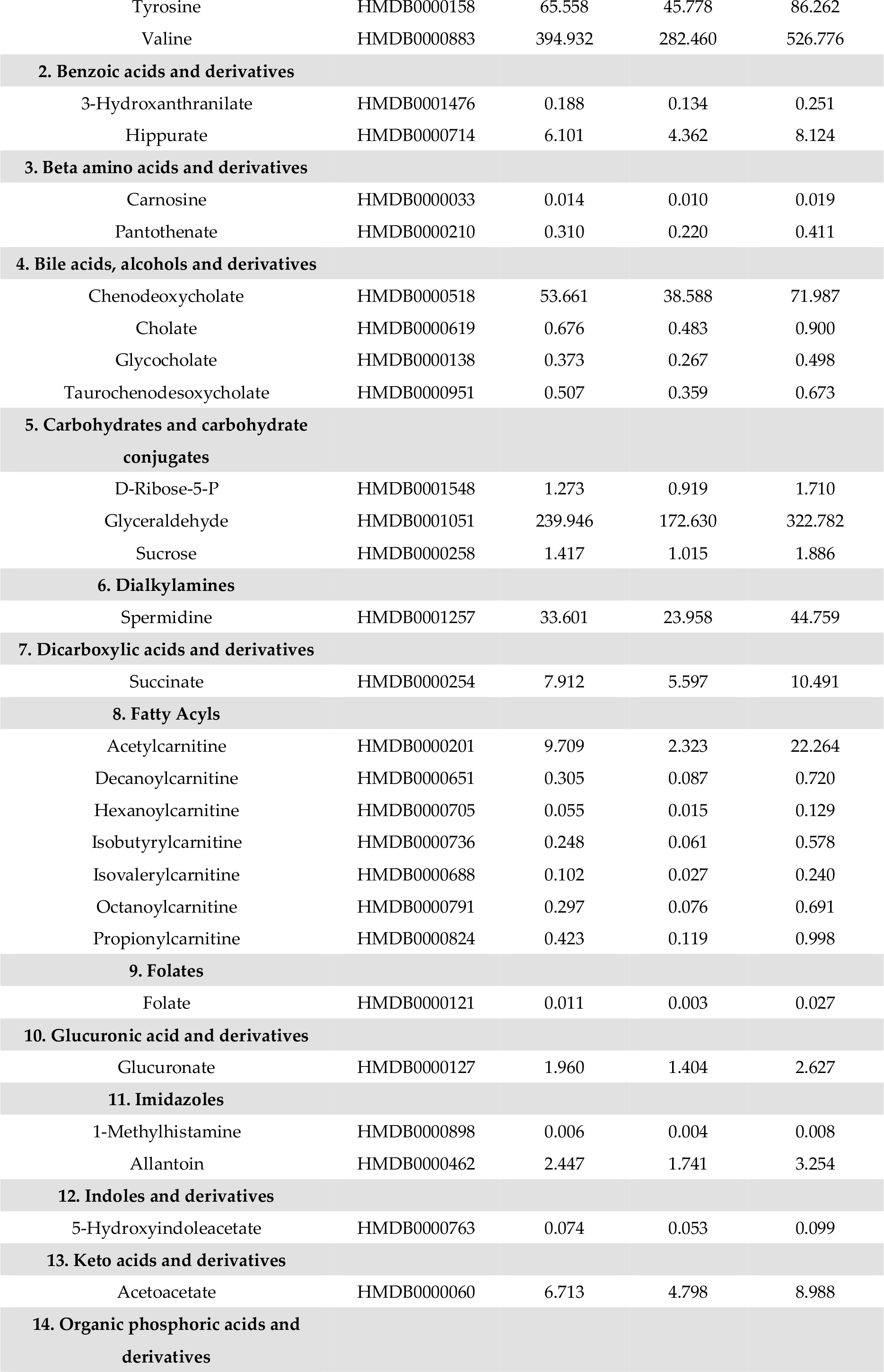

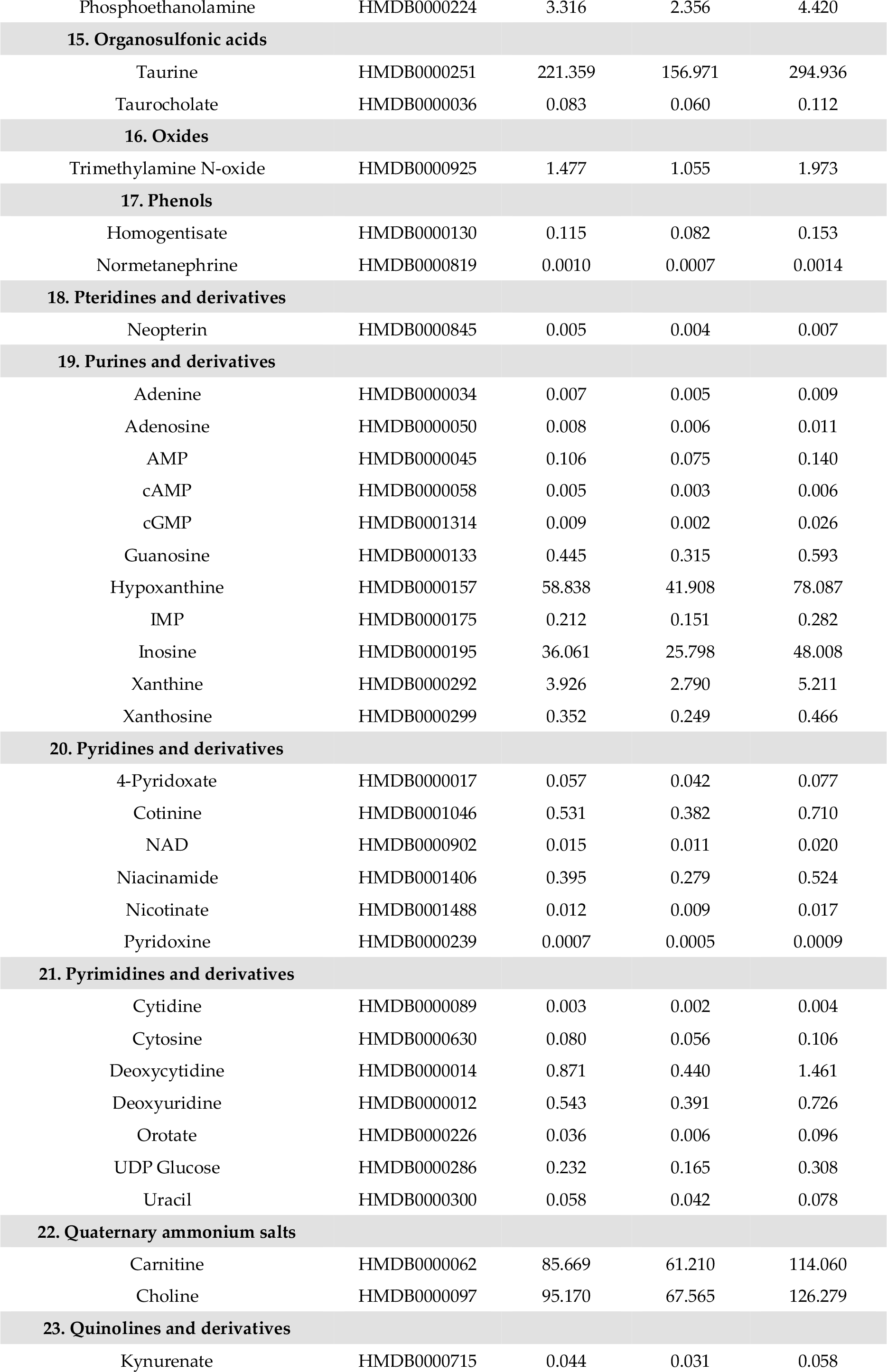

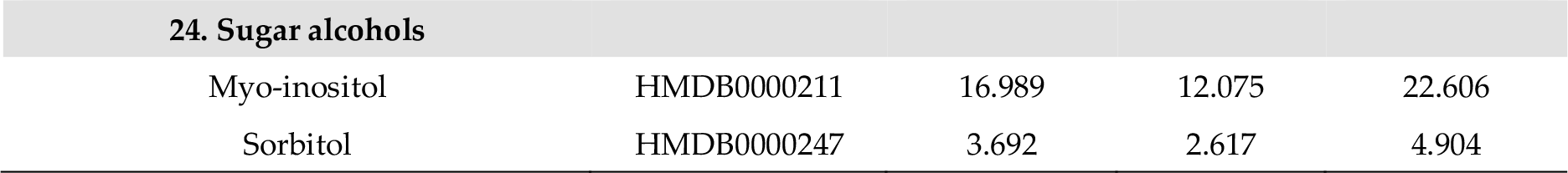

#### 3.2.8 Robustness and cross-platform comparison

To verify the performance of our method, we analyzed the NIST standard reference material SRM 1950 plasma. The correlation coefficient for 17 matched metabolites between the given reference values and from our semi-quantitative method was 0.967, indicating the high performance of our method (Figure 10A).

Furthermore, we verified cross-platform comparability. This was achieved by comparing metabolite concentrations analyzed using our method against two completely different analytical platforms (BIOCRATES AbsoluteIDQ p180 kit and NMR) in our QC samples. We obtained a high correlation coefficient for matched 38 metabolites measured using BIOCRATES AbsoluteIDQ p180 kit and our method (R^2^=0.975) (Figure 10B) and for matched 22 metabolites measured using NMR and our method (R^2^=0.884) (Figure 10C). These results demonstrate the robustness of our method.

**Figure 10.**
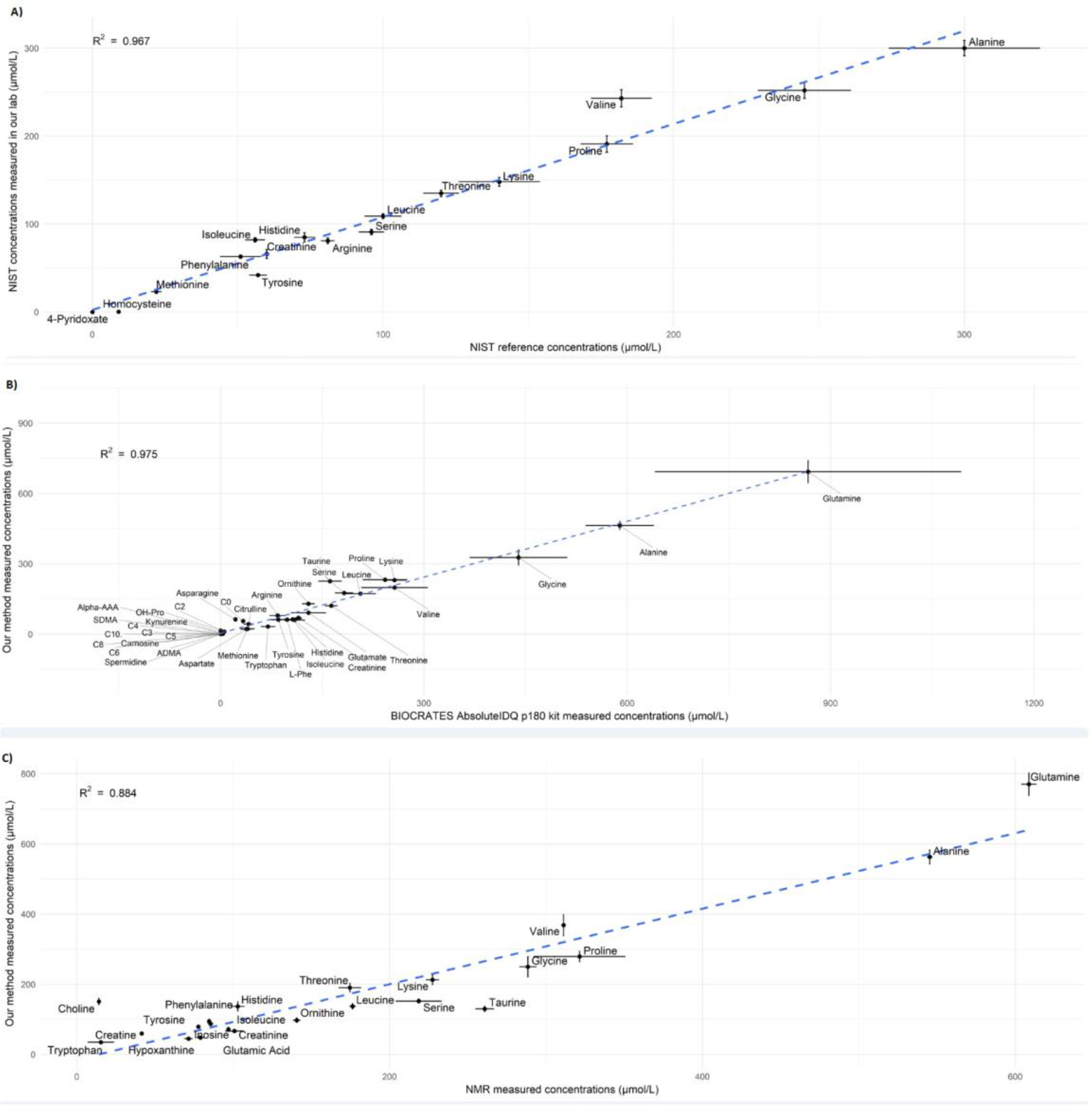
(A) Comparison of metabolite concentration reference values given in the NIST SRM 1950 plasma against our method. Comparison of metabolite concentrations in our QC samples measured between our method and (B) BIOCRATES AbsoluteIDQ p180 kit, and (C) NMR analyses. Error bars represent 95% confidence intervals.

### Automated data processing

After the raw data processing using the instrument-coupled software (TargetLynx), a single-sheet excel file containing information such as sample ids, file name, and concentrations (PPB) is generated in a complex format for all metabolites. For example, in a single-batch run of 85 samples, after data processing the concentration values for all samples are obtained under each metabolite separately. This means that if a data matrix of 85 samples x 100 metabolites is desired, each individual concentration value must be copied and pasted in another sheet. PPB values are then converted into μM and corrected for process efficiencies and dilution factors (if any) and normalized with tissue weight or cell number depending on the sample type. Apart from these steps, as the samples are analyzed in a randomized manner, the next step is to rearrange the experimental samples according to phenotypic group and also to separate the QC samples. Manual data formatting to create a ready-to-use data matrix for visualization and for downstream statistical analyses is a tedious and time-consuming task and (most importantly) is prone to errors.

We have implemented a software package “Unlynx” in R statistical language. This package takes the raw data produced by TargetLynx software as input and produces processed data into ready-to-use spreadsheets. Using the R package, all the PPB values are converted into μmol/L, μmol/g, or μmol/million cells by diving with molecular weight of the respective metabolites and correction factor (weight of tissues, number of cells, and dilution factor) with respect to sample types. In addition, mean concentrations and %CV of QC samples for all the metabolites are calculated and used to evaluate quality by comparing with the *in-house* QC database constructed based on inter-day %CV. Furthermore, retrieving LLOQ, ULOQ, and outlier values in each phenotypic group according to one and two standard deviations, retention time values, and R^2^ values for each metabolite is implemented in the R software. After implementing the automated data processing, we had reduced the 2-day manual workload to a few minutes.

Thus, after processing the data using TargetLynx, we routinely export all results to an excel file for automated data processing using “Unlynx” software. Typically, the data resulting from such automated data processing is suitable for more specialized data analyses (such as statistical hypotheses testing, classification, regression, and clustering) aimed at answering specific scientific questions related to the study design.

### Applicability of the method

We have applied our fully validated analytical methodology in various international and national biomedical research projects, epidemiological studies, clinical studies including dietary interventions, and clinical trials. We have successfully implemented our technology in the following research fields, including but not limited to: mitochondrial metabolism/disorders [11,17–23], cancer [24,25], bone metabolism [12], endocrinology [26,27], psychiatric disorders [28], inflammatory bowel disease [29], viral infections [30–33], allergies [34,35], circadian rhythms [36], and pain research [37].

## Conclusions

The developed high-throughput targeted and semi-quantitative method was optimized for various biological matrices (biofluids, tissues, cells) from different organisms. We validated the analytical method according to EMA guidelines for bioanalytical methods and showed good accuracy, reproducibility, selectivity, specificity, recoveries, and stability. We have also implemented a strict quality management and electronic notebook system. Reproducibility was demonstrated by consistent results for retention time and correlation coefficient of calibration curves, and concentrations of QC samples over a period of 1 year. Reliability was shown by the excellent correlation between metabolite concentrations measured using our method and the NIST reference values. Moreover, robustness was shown via good cross-platform comparability between two completely different analytical platforms. Furthermore, we have automated the downstream data processing steps to handle sample analyses in a high-throughput manner, which is particularly valuable for analyzing population cohorts and large clinical samples for metabolomics studies. We have successfully applied this method in many biomedical research projects and clinical trials, including epidemiological studies for biomarker discovery.

## Supplementary Materials

The following are available online at www.mdpi.com/link, Table S1: List of metabolites showing molecular weight, retention time, linearity of calibration, and compound dependent MS parameters.

## Acknowledgments

The authors would like to thank all the collaborators, especially Prof. Anu Wartiovaara for providing the biological samples; Jean-Christophe Yorke, Vasudev Kantae, Kenneth Nazir, and Bharat Gajera for technical assistance, and Dr. Janne Wallenius for reviewing the manuscript. The Biocenter Finland and HiLIFE supported this work.

## Author Contributions

J.N. performed all the validation experiments, processed the data, and wrote the manuscript. M.K. and J.N. were involved in extraction protocol optimization. G.P. developed the R package (Unlynx) for automated data processing. A.P. performed the QC data analysis, prepared the related table, and wrote the corresponding section. V.V. conceived, designed, and supervised the study, and wrote and edited the manuscript. All authors edited the manuscript.

## Conflicts of Interest

The authors declare no conflict of interest. The sponsors had no role in the design of the study; in the collection, analyses, or interpretation of data; in the writing of the manuscript; and in the decision to publish the results.

